# Idiosyncratic variation in the fitness costs of tetracycline-resistance mutations in *Escherichia coli*

**DOI:** 10.1101/2020.09.12.294355

**Authors:** Kyle J. Card, Jalin A. Jordan, Richard E. Lenski

## Abstract

A bacterium’s fitness relative to its competitors, both in the presence and absence of antibiotics, plays a key role in its ecological success and clinical impact. In this study, we examine whether tetracycline-resistant mutants are less fit in the absence of the drug than their sensitive parents, and whether the fitness cost of resistance is constant or variable across independently derived lines. Tetracycline-resistant lines suffered, on average, a reduction in fitness of almost 8%. There was substantial among-line variation in the fitness cost. This variation was not associated with the level of phenotypic resistance conferred by the mutations, nor did it vary significantly across several different genetic backgrounds. The two resistant lines with the most extreme fitness costs involved functionally unrelated mutations on different genetic backgrounds. However, there was also significant variation in the fitness costs for mutations affecting the same pathway and even different alleles of the same gene. Our findings demonstrate that the fitness costs of antibiotic resistance do not always correlate with the phenotypic level of resistance or the underlying genetic changes. Instead, these costs reflect the idiosyncratic effects of particular resistance mutations and the genetic backgrounds in which they occur.

## Introduction

Antibiotics are an essential component of modern medicine. Although they have dramatically reduced the morbidity and mortality caused by severe bacterial infections, their benefits have diminished in recent years because of their overuse in the clinic and in agriculture, which has led to the evolution and proliferation of antibiotic-resistant pathogens. As a result, many infections have become more difficult to treat with mainline drug therapies, and in severe cases, some pathogenic strains have become resistant to all available drugs. An understanding of the forces underlying and shaping antibiotic resistance is therefore critical to the future health of the human population.

Bacteria can evolve resistance by either spontaneous mutations or horizontal acquisition of resistance genes. Spontaneous mutations commonly confer resistance by altering the cellular target of the antibiotic or increasing its efflux (Blair et al. 2015). Mechanisms associated with horizontal gene transfer include target modification, drug detoxification, and the acquisition of novel efflux pumps (Blair et al. 2015). In either case, resistant variants have a clear advantage over their sensitive counterparts when exposed to the corresponding antibiotic. However, these resistant types often suffer fitness costs because they disrupt the normal functioning of metabolic pathways and physiological processes or increase the energetic burden on the cell (Lenski and Bouma 1987; Nguyen et al. 1989; Andersson and Hughes 2010; Vogwill and MacLean 2015). Resistant types should therefore have lower growth rates than, and be outcompeted by, their sensitive counterparts in the absence of drugs.

A resistant bacterium’s competitive fitness, both in the presence and absence of a drug, is an important factor that contributes to its ecological success and thus its clinical impact (Lenski 1997; Vogwill and MacLean 2015; Hughes and Andersson 2017). For example, the fitness of a resistance mutation determines its likelihood of persisting in a bacterial population prior to drug exposure, its maintenance in a population at a particular drug concentration, and its reversibility when the antibiotic is reduced or removed from the environment (Hughes and Andersson 2017; Santos-Lopez et al. 2019).

The expected time required to reduce the frequency of a resistant mutant in a bacterial population following the cessation of antibiotic use is inversely proportional to the fitness cost of the resistance mutation (Lenski 1997). Although mathematical models can predict the rate of these frequency declines (Levin et al. 1997), these theoretical expectations often are not met under real-world scenarios for at least two reasons. First, some resistance mechanisms are inherently cost free, at least in certain environments. Several mutations in the gene *rpsL* confer resistance to streptomycin, but they have little or no fitness cost in both *Escherichia coli* and *Salmonella typhimurium* (Tubulekas and Hughes 1993), and they even confer a competitive advantage over wild-type strains in some animal infection models (Björkman et al. 1998; Enne et al. 2005). These cost-free *rpsL* mutations are also found in streptomycin-resistant *Mycobacterium tuberculosis* populations, where they may facilitate the long-term maintenance of this resistant type (Böttger et al. 1998; Andersson and Hughes 2010). Similarly, treatment of *Helicobacter pylori* infections with clarithromycin has been found to select for highly resistant commensal *Enterococcus* species that persist for years after drug treatment (Sjölund et al. 2003). This last outcome demonstrates a troubling side-effect of antibiotic use, in which the microbiome can act as both a reservoir for resistance genes and as a conduit for their horizontal transfer to pathogens (Sommer et al. 2010).

Second, pleiotropic costs associated with chromosomal- or plasmid-mediated resistance can often be reduced or even eliminated through subsequent compensatory evolution (Bouma and Lenski 1988; Schrag et al. 1997; Kugelberg et al. 2005; Nilsson et al. 2006; Andersson and Hughes 2010; Barrick et al. 2010). For example, clinically relevant levels of fluoroquinolone resistance occur through the sequential substitution of mutations in several genes (Lindgren et al. 2003). Early genetic changes in the mutational pathway exact a cost on bacterial growth in both laboratory media and mouse models, but the cost can be ameliorated through later resistance mutations (Marcusson et al. 2009). Thus, evolution can restore a bacterial population’s ancestral growth rate in the absence of drug selection while simultaneously preserving resistance in the event of future exposure to antibiotics. Moreover, compensatory evolution can sometimes drive multidrug resistance; this outcome has been seen when a genetic change simultaneously provides resistance to a newly imposed drug while reducing the fitness cost associated with resistance to a previous antibiotic (Trindade et al. 2009). Compensatory evolution shows how pleiotropic effects of one mutation can set the stage for epistatic interactions with subsequent mutations.

In general, a bacterium’s genetic background can influence the fitness costs of antibiotic resistance. For example, Vogwill and colleagues (2016) examined the costs of rifampicin-resistance mutations in the gene *rpoB* across several *Pseudomonas* species. They found that some mutations vary in their fitness effects across backgrounds, and these costs correlate with transcriptional efficiency. Thus, the same *rpoB* mutation can differentially affect transcriptional efficiency depending on the genetic background, and these idiosyncratic effects in turn lead to heterogeneity in costs. This work evaluated genetic-background effects across a fairly broad phylogenetic scale, while focusing on mutations in a single gene. One can also ask whether genetic background affects the fitness cost of resistance even among recently diverged clones of a single species, and for resistance that has evolved through more diverse mutational pathways.

To address these issues, we evaluated the competitive fitness in the absence of drugs of tetracycline-resistant clones that evolved from several different *E. coli* backgrounds, which previously diverged during a long-term evolution experiment (LTEE). We ask several questions. First, is there a fitness cost to resistance? Second, is the cost greater for mutants that evolved higher levels of resistance (Fig. 1A)? Third, do fitness costs vary in an idiosyncratic manner that does not depend on the level of resistance achieved (Fig. 1B)? Fourth, if there is indeed idiosyncratic variation among lines in the cost of resistance, what factors contribute to that variability? On balance, we found that the resistant lines are indeed less fit than their sensitive counterparts. These fitness costs do not correlate with the level of resistance achieved, nor do they vary among the several genetic backgrounds that we examined (Card et al. 2019). Some variation in cost of resistance occurs even among different mutations in the same gene, on the same genetic background, and conferring the same phenotypic resistance. In any case, further research on the fitness effects of antibiotic resistance should be pursued because of its potential implications for public health and patient treatment.

**Figure 1.**
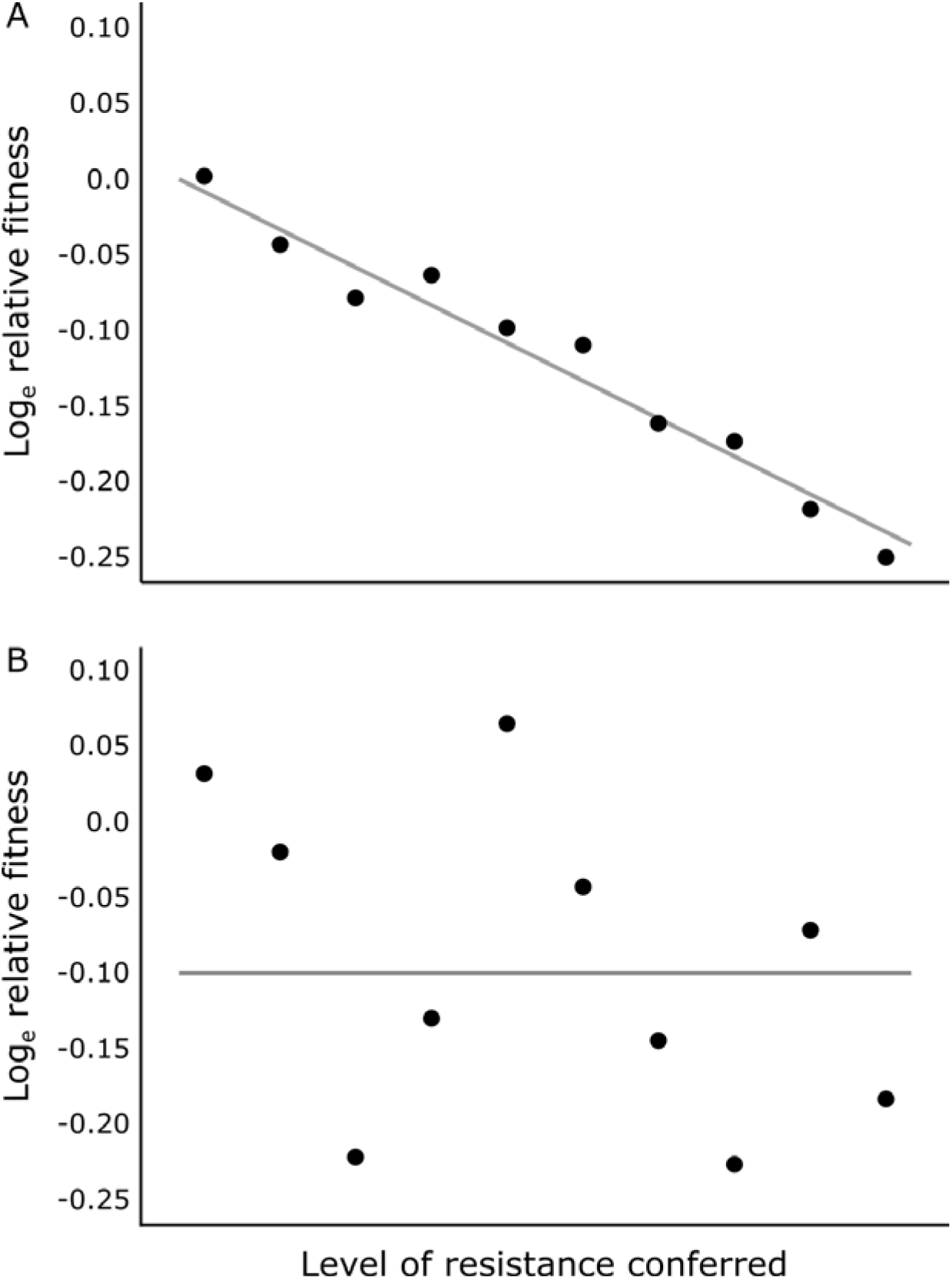
Schematic illustration of fitness effects of antibiotic resistance mutations under two scenarios. (A) Tradeoff model, in which the fitness of a resistant line, when measured in the absence of drugs, is negatively correlated with the level of resistance conferred by its mutations. (B) Idiosyncratic model, in which the fitness of resistant lines varies for reasons unrelated to the level of resistance. This idiosyncratic variation might, in principle, reflect differences between genetic backgrounds, mutations in different target genes, different alleles of the same target gene, secondary mutations, and epistatic interactions between the resistance mutations and their genetic backgrounds. The fitness of each resistant line is expressed relative to its sensitive counterpart. A log-transformed relative fitness of 0 indicates no fitness cost associated with resistance, while values below and above 0 represent fitness costs and benefits, respectively.

## Materials and Methods

### EXPERIMENTAL CONDITIONS AND BACTERIAL STRAINS

The LTEE has been described in detail elsewhere (Lenski et al. 1991; Lenski 2017). In brief, 12 replicate populations of *E. coli* were founded from a common ancestral strain, called REL606 (Daegelen et al. 2009). These populations have been propagated for over 32 years and 73,000 generations by daily 100-fold dilutions in Davis Mingioli minimal medium supplemented with 25 μg/mL glucose (DM25).

In this study, we examined the competitive fitness of tetracycline-resistant mutants that evolved from the LTEE ancestor and clones sampled from four LTEE populations (denoted Ara−5, Ara−6, Ara+4, and Ara+5) after 50,000 generations. Specifically, we analyzed 4 mutants that independently evolved from the ancestral background, and 3 mutants that evolved from each derived background, for a total of 16 mutants (Table S1). We also used three clones as common competitors: REL607, REL10948, and REL11638. REL607 is a spontaneous Ara^+^ mutant of REL606, the LTEE ancestor (Lenski et al. 1991). REL10948 is an Ara^−^ clone isolated from the Ara−5 population at 40,000 generations, and REL11638 is a spontaneous Ara^+^ mutant of that clone (Wiser et al. 2013; Lenski et al. 2015). The Ara marker is selectively neutral in the glucose-limited medium; it serves to differentiate competitors during fitness assays because the Ara^−^ and Ara^+^ cells form red and white colonies, respectively, on tetrazolium-arabinose (TA) agar. We used REL607 as the common competitor for REL606 and the four tetracycline-resistant clones derived from it. The 40,000-generation clones served as common competitors for the four 50,000-generation parental clones and twelve resistant mutants that evolved from them; using these common competitors ensured that the differences in fitness were not so large that their densities would fall below the detection limit during the fitness assays.

### FITNESS ASSAYS

Assays were performed in the absence of antibiotics to assess the relative fitness of drug-resistant mutants and their susceptible counterparts. Fitness was measured in an environment identical to that of the LTEE, except the medium contained 250 μg/mL glucose (DM250). Resistant mutants and their sensitive parents each competed, in paired assays, against the same common competitor with the opposite Ara-marker state (Fig. 2). To set up each competition assay, the competitors were revived from frozen stocks, and they were separately acclimated to the culture medium and other conditions over two days. The competitors were then each diluted 1:200 into fresh medium, and a sample was immediately plated on TA agar to assess their initial densities based on colony counts. The competition cultures were then propagated for 3 days, with 1:100 dilutions each day in fresh medium. At the end of day 3, a sample was plated on TA agar to assess the competitors’ final densities. We quantified the realized growth rate of each competitor based on its initial and final density and the net dilutions imposed (Lenski et al. 1991). We then calculated relative fitness as the ratio of the realized growth rate of the clone of interest (either a resistant clone or its sensitive parent) to that of the common competitor. Lastly, the fitness of a resistant mutant in each assay was normalized by dividing it by the relative fitness of the paired assay obtained for its parental strain. We performed a total of 80 pairs of fitness assays (160 competitions in total) to produce 5 replicate estimates of the fitness of each of the 16 tetracycline-resistant mutants relative to its sensitive parent. The relative fitness values were log_e_-transformed before the statistical analyses reported in the Results below.

**Figure 2.**
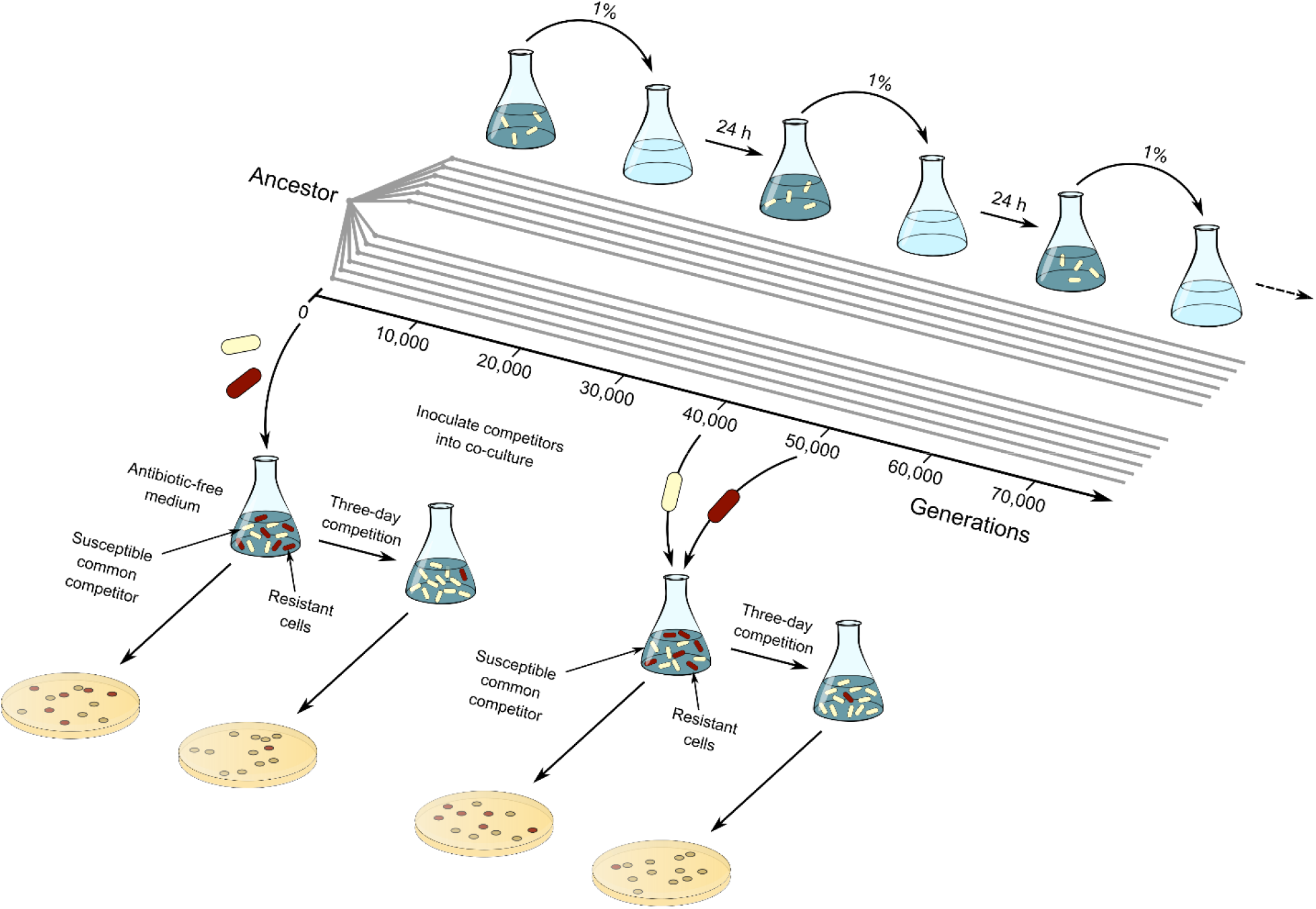
Schematic illustration showing the derivation of the strains used in this study and the methods employed to measure the fitness of resistant lines relative to their sensitive parents. Twelve initially identical *E. coli* populations were founded from the same ancestral strain to start the LTEE. A genetic marker distinguishes two sets of six populations each. These populations have evolved for >73,000 generations with daily transfers in a minimal glucose medium. In paired assays, we examined the fitness of tetracycline-resistant mutants (shown in red) that evolved either from the LTEE ancestor or one of four clones sampled at generation 50,000 by competing them against marked susceptible competitors (shown in yellow). We used REL607 as the common competitor for the LTEE ancestor and resistant lines evolved from it, and two 40,000-generation clones (see Materials and Methods) as common competitors for the derived parental strains and their evolved resistant lines. After acclimation to the culture conditions, competitors were mixed at an equal volumetric ratio in a common medium. These cultures were propagated for three days in the absence of tetracycline by serial 1:100 transfers. We quantified each competitor’s realized growth rate from the initial and final densities after plating on TA agar, taking into account the net dilution over the three days. These realized growth rates were then used to calculate the fitness of a resistant line relative to its sensitive parent (see Materials and Methods).

## Results

### TETRACYCLINE-RESISTANT LINES HAVE REDUCED FITNESS IN THE ABSENCE OF THE ANTIBIOTIC

We ask first whether tetracycline resistance is costly, on average, in the absence of the drug. The grand mean of the log_e_-transformed fitness of the 16 resistant lines relative to their paired parental strains is −0.0771, indicating that the resistant mutants grow ~7.7% more slowly than their sensitive counterparts during head-to-head competitions with a common competitor. This value differs significantly from the null hypothesis that the resistant lines and their sensitive parents are equally fit (*t_s_* = 2.9973, 15 d.f., one-tailed *p* = 0.0045).

### COST OF RESISTANCE VARIES AMONG RESISTANT MUTANTS

We measured the relative fitness of each resistant line with 5-fold replication. This replication allows us to test whether the variation in fitness among the 16 tetracycline-resistant lines is simply measurement noise or, alternatively, reflects genetic variation in the cost of resistance. Table 1 shows the analysis of variance (ANOVA). The variation among the 16 lines is about 10-fold greater than expected from the variation between replicate assays performed on the same line (*F*_15,64_ = 10.34, *p* ≪ 0.0001).

**Table 1.** ANOVA on the log-transformed fitness estimates of 16 tetracycline-resistant lines, each measured relative to its sensitive parent.

| Source | SS | d.f. | MS | *F* | *p* |
| --- | --- | --- | --- | --- | --- |
| **Line** | 0.7948 | 15 | 0.0530 | 10.3384 | << 0.0001 |
| **Error** | 0.3280 | 64 | 0.0051 |  |  |
| **Total** | 1.1228 | 79 |  |  |  |

There are many possible reasons why the cost of resistance might vary including mutations in different genes, different alleles even of the same gene, different genetic backgrounds, epistatic interactions between mutations and genetic backgrounds, and so on. In the sections that follow, we examine various possibilities.

### POSSIBLE REVERSIONS OF UNSTABLE MUTATIONS DO NOT EXPLAIN THE VARIATION IN FITNESS COST

We previously sequenced the complete genomes of the 16 resistant lines, and we compared them to their parental strains to identify the mutations responsible for their resistance (Card et al. 2020). Two lines had no identifiable mutations (Ara−5-1 and Ara+5-1), even though they had increased phenotypic resistance relative to their respective parent strains (Card et al. 2019). This discrepancy suggested that these two resistant lines may have had unstable genetic changes, which might have reverted prior to the genomic analysis and our fitness assays. Potentially unstable mutations include changes in the copy number of oligonucleotide repeats and gene amplifications. We repeated the ANOVA, except excluding the two resistant lines without identifiable mutations. The variation in the cost of resistance remains highly significant in the 14 lines with known, stable mutations (*F*_13,56_ = 10.15, *p* ≪ 0.0001).

### LEVEL OF PHENOTYPIC RESISTANCE DOES NOT EXPLAIN THE VARIATION IN FITNESS COST

All of the resistant lines evolved during a single round of exposure to tetracycline. However, they vary in the resulting minimum inhibitory concentration (MIC) that they achieved. They also vary in the magnitude of the increase in their MICs relative to their parental strains, which also varied in their MICs. It is possible that mutations that provide greater resistance have higher fitness costs (Fig. 1A). To test that possibility, we examined the correlation between the log-transformed fitness values of the 14 resistant lines and their log-transformed MICs, as previously reported (Card et al. 2019). However, the correlation is not significant; in fact, it is weakly positive (*r* = 0.1682, two-tailed *p* = 0.5655). We also computed the correlation between the log-transformed fitnesses and log-transformed fold-increases in resistance, but again the correlation is weakly positive and not significant (*r* = 0.1002, two-tailed *p* = 0.7332). Thus, we find no evidence that the variation in the fitness cost of tetracycline resistance is related to the level of resistance conferred by the underlying mutations.

### GENETIC BACKGROUND DOES NOT EXPLAIN THE VARIATION IN FITNESS COST

The 14 tetracycline-resistant mutants with identifiable mutations evolved on five different genetic backgrounds. We asked whether the average cost of resistance differed between the backgrounds. In this case, the ANOVA tests whether the variance in the average cost of resistance for mutants derived from different backgrounds is greater than expected given the variance in the average cost for mutants derived from the same background. This analysis indicates no significant effect of the genetic background on the cost of resistance (*F*_4,9_ = 0.47, *p* = 0.7570).

### IDIOSYNCRATIC DIFFERENCES BETWEEN MUTANT LINES IN THE COST OF RESISTANCE

Neither the level of phenotypic resistance conferred by mutations nor the genetic background in which they arose explains the substantial variation in the fitness effects of tetracycline resistance. Instead, it appears there are idiosyncratic differences in the fitness costs associated with different resistance mutations (Fig. 1B). These idiosyncratic effects could, in principle, reflect mutations in different genes, different mutations in the same target gene, secondary mutations that might have hitchhiked with the mutations conferring resistance, or epistatic interactions between any of these new mutations and the existing mutations that distinguished the different parental strains. Without a much larger number of resistant lines, it is not possible to rigorously disentangle these various sources of idiosyncratic fitness costs. However, by examining and contrasting specific cases, we are able to shed light on some of the sources of these differences.

Two resistant clones, Ara+4-3 and Ara+5-2, have fitness costs that are very similar to one another, but more than double the cost of any of the other 12 resistant mutants (Fig. 3). Yet these two cases occurred on different genetic backgrounds and have different mutations. Ara+4-3 has mutations in *hns*, which encodes a histone-like global regulator, and *lpcA*, which encodes a phosphoheptose isomerase; Ara+5-2 has a single mutation in *ompF*, which encodes an outer-membrane porin (Card et al. 2020). We asked whether these two extreme cases are solely responsible for the heterogeneity in fitness costs by performing an ANOVA that excludes them. The variation in fitness costs among the other 12 clones is reduced, but it nonetheless remains highly significant (*F*_11,48_ = 4.44, *p* = 0.0001).

**Figure 3.**
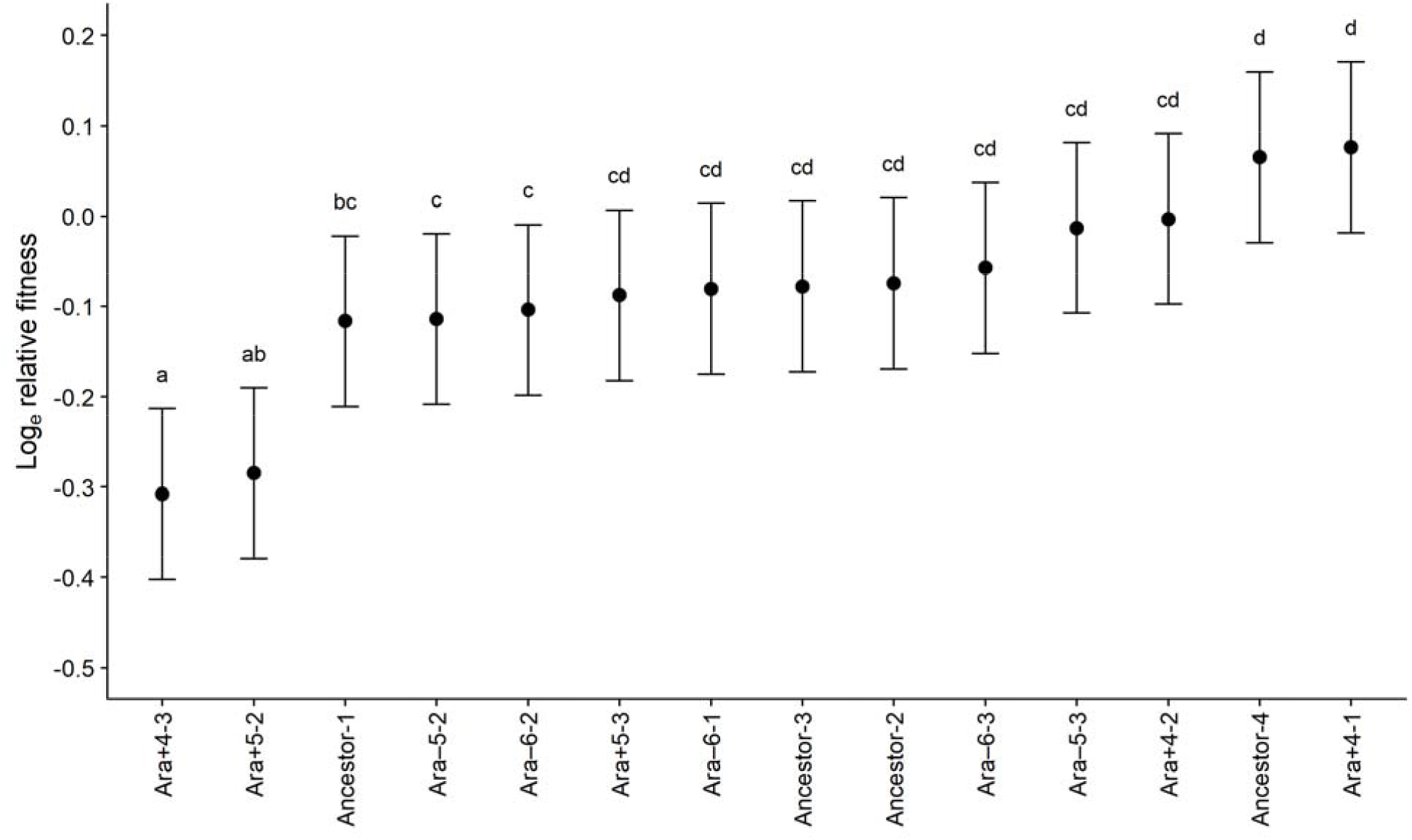
Fitnesses of 14 tetracycline-resistant mutants relative to their parental strains. The mutants are arranged from lowest to highest fitness. Each symbol shows the mean log_e_-transformed fitness, based on 5-fold replication of paired fitness assays. Error bars show 95% confidence limits calculated using the *t*-distribution with 4 d.f. and the pooled standard deviation estimated from the ANOVA (Table 1). Letters above the error bars identify mutants with relative fitnesses that are not significantly different, based on Tukey’s “honestly significant difference” test for multiple comparisons.

Nine of the 14 resistant clones have a single mutation each, while four of them (Ara−5-2, Ara−6-2, Ara+4-3, Ara+5-3) have two mutations, and another (Ancestor-2) has three mutations (Card et al. 2020). It is reasonable to imagine that in each clone one mutation confers the drug resistance, while the others merely hitchhiked with the resistance mutation. Such hitchhikers might include deleterious mutations that reduce fitness. Therefore, we compared the fitness costs for the resistant clones with and without secondary mutations. The average fitness cost of the clones with multiple mutations is higher (13.8%) than the average of those with single mutations (5.5%), but the difference is only marginally nonsignificant given the small number of clones in each group and the high variation within each group (Welch’s *t*-test, *t_s_* = 1.4751, 9.3 d.f., one-tailed *p* = 0.0866).

It is also interesting to compare the four resistant clones derived from the ancestral LTEE background. All four resistant clones evolved the same level of phenotypic resistance, with MICs that are 4-fold higher than their parental strain (Card et al. 2019). Moreover, all four have mutations affecting the same two-component system that regulates the synthesis of outer-membrane proteins: one clone (Ancestor-1) has a 11-bp deletion in *envZ*, which encodes the sensory histidine kinase; the others (Ancestor-2, Ancestor-3, Ancestor-4) have nonsynonymous mutations in *ompR*, which encodes the DNA-binding response regulator. Even with these striking phenotypic and genetic similarities, an ANOVA shows significant heterogeneity in the fitness of these clones (*F*_3,16_ = 4.50, *p* = 0.0180). We can also compare only Ancestor-3 and Ancestor-4 (each having a single mutation in *ompR* and no other mutation), and the variation in fitness remains significant (*F*_1,8_ = 5.71, *p* = 0.0439). These results show that different mutations in the same target pathway, and even different alleles of the same gene, can lead to different fitness costs of drug resistance.

## Discussion

In previous work, we examined the role that genetic background plays in both the phenotypic and genotypic evolution of antibiotic resistance. First, we examined the potential of several different LTEE backgrounds to evolve increased resistance to several antibiotics. We found that evolvability was idiosyncratic with respect to the parental genotype, such that resistance was more constrained in some backgrounds than in others (Card et al., 2019). Genetic differences will accumulate between populations, even if they evolve in the same permissive environment. These differences can unpredictably alter their ability to respond evolutionarily when challenged with antibiotics. Second, we sequenced the complete genomes of some of these resistant mutants and assessed whether the different initial genotypes took similar or divergent mutational paths to increased resistance (Card et al. 2020). Again, we found that the initial genetic background is important. On average, the replicate lines that evolved from the same founding genotypes had more gene-level mutations in common than lines derived from different founding genotypes.

The aim of this study was to examine whether and how genetic background influences the fitness effects of resistance mutations in the absence of antibiotic. In particular, we examined the fitness costs of tetracycline resistance in 16 lines that evolved from five sensitive parental backgrounds. We found that the resistant lines are, on average, less fit than their sensitive counterparts in the absence of the antibiotic. This result is not surprising, given that resistance mutations often disrupt the normal function of metabolic or physiological processes, or impose energetic demands that reduce growth and competitiveness (Andersson and Hughes 2010). We also observed highly significant variation among the resistant lines in their fitness costs (Table 1). This variation remained substantial (Fig. 3) even after we excluded two strains without identified mutations (Card et al. 2020). These two strains exhibited phenotypic resistance in our earlier work (Card et al. 2019), but that resistance might have been conferred by unstable genomic changes, such as gene amplifications or frameshift mutations in homopolymeric tracts that can cause “phase variation” (Moxon et al. 1994). If so, these unstable changes could have reverted prior to the genomic analysis and the competition assays that we performed.

We then addressed two broad possibilities regarding the variation in fitness cost between the 14 lines with known, stable mutations. First, we asked whether there is a relation between a line’s phenotypic resistance and its fitness cost, such that mutations that confer greater resistance are more costly (Fig. 1A). A meta-analysis of fitness costs across several species and drug classes by Melnyk and colleagues (2015) supported this association, and the authors suggested it could be understood from evolutionary and mechanistic perspectives. Imagine a population that is well-adapted to one environment and hence near a local fitness optimum. If the environment changes, such as with the addition of an antibiotic, then the population may evolve toward a different optimum through the substitution of new mutations. Mutations of large effect will bring the population closer to this new optimum than mutations of small effect. However, if the environment later reverts to its previous state, then populations that substituted the large-effect mutations will be further from their previous optimum than those populations that acquired small-effect mutations. From a mechanistic standpoint, the increased expression of efflux pumps or drug targets diverts resources from other cellular processes. Also, resistance mutations that change evolutionarily conserved proteins are more likely to disrupt their functions than improve them. In our study, however, there was no significant association between fitness costs and the level of resistance conferred by mutations, whether on an absolute basis or relative to the parent strain.

The second broad possibility is that the fitness costs of resistance can vary for reasons unrelated to the level of resistance conferred (Fig. 1B). There are several potential reasons for such idiosyncratic variation. One possibility is that the same resistance mutation may have different fitness costs in different genetic backgrounds. In *Campylobacter jejuni*, for example, a C257T mutation in the gene *gyrA* confers fluroquinolone resistance. When fluroquinolone-resistant and -susceptible strains were inoculated separately into chickens, they colonized equally well and each persisted even in the absence of drug exposure (Luo et al. 2005). However, when resistant and sensitive strains were co-inoculated, the resistant variants often prevailed. Further work indicated that this particular *gyrA* mutation was beneficial in some genetic backgrounds, even in the absence of antibiotic, and costly in others (Luo et al. 2005). In our study, by contrast, the variation in fitness costs among strains was not explained by genetic-background effects, but instead involved several other factors.

One such factor is that resistance mutations can occur in different genes, which can lead to different fitness costs. In this study, the relative fitnesses of clones Ara+4-3 and Ara+5-2 were significantly lower than the other 12 strains. Ara+4-3 is the only line with mutations in either *lpcA* or *hns*. Mutations in the former gene have been shown to confer tigecycline resistance in *E. coli* through modifications to the lipopolysaccharide biosynthesis pathway, and these mutations have moderate fitness costs in vitro (Linkevicius et al. 2013, 2016). The latter gene encodes the global transcriptional regulator H-NS, and mutations in it affect acid resistance (Giangrossi et al. 2005), the modulation of osmotic stress (Lucht et al. 1994), and several other important cellular processes. Changes to this regulator’s structure and function might therefore have large fitness costs via widespread pleiotropic effects. The Ara+5-2 clone evolved a 9-bp insertion in *ompF*, which encodes the sole major porin in the LTEE ancestral strain (Crozat et al. 2011); this mutation presumably reduces the cell’s antibiotic uptake, but at the expense of acquiring nutrients (Ferenci 2005; Phan and Ferenci 2017). Thus, resistance mutations that affect different cellular pathways and functions can have variable fitness costs, a finding that is consistent with many other studies (Vogwill and MacLean 2015).

Another factor is that mutations in different genes that are part of the same physiological pathway may confer similar resistance levels but have different fitness costs. In our study, four tetracycline-resistant lines derive from the same LTEE ancestor: one had a mutation in *envZ*, while the other three had mutations in *ompR*. These genes encode proteins that comprise a two-component regulatory system that regulates cellular responses to osmotic stress, and which affects antibiotic resistance through altered expression of the major porin OmpF (Chakraborty and Kenney 2018; Choi and Lee 2019). We observed significant heterogeneity in fitness even among these lines, implying that different changes within this one pathway can impose unique burdens. The evolution of carbapenem resistance in *E. coli* K12 can also occur by mutations in this same two-component system, again with variable fitness costs (Adler et al. 2013). In their study, Adler and colleagues (2013) found that *envZ* mutants had no measurable loss of fitness in the absence of antibiotic, whereas *ompR* mutations suffered a large cost. By contrast, in our study the *envZ* mutation was more costly, which may reflect differences between the *E. coli* K12 and B strain backgrounds or the use of different culture media.

Yet another factor is that different mutations in the same gene can have different costs. The evolution of rifampicin resistance, for example, typically occurs via mutations in several canonical regions of *rpoB*, which encodes the β subunit of the RNA polymerase (Reynolds 2000; Ahmad et al. 2002; Barrick et al. 2010; MacLean et al. 2010; Hall and MacLean 2011). Different alleles have widely varying costs that impact their competitive ability and, moreover, affect the dynamics of subsequent compensatory evolution (Barrick et al. 2010). In our study, two clones derived from the same parent had different nonsynonymous mutations in *ompR*. Both conferred the same level of resistance to tetracycline, but they had different fitness costs in the absence of the drug. Such differences can have important public-health consequences, because a resistant lineage’s competitive fitness in the absence of antibiotics is critically important for its long-term persistence in a heterogeneous environment.

More generally, we argue that further studies of the fitness costs of antibiotic resistance are needed, because this phenomenon can inform treatment strategies. Standard clinical practice calls for aggressive treatment to eliminate an infecting pathogen before it has time to evolve resistance (Craig 2001; Drlica 2003; Mehrotra et al. 2004; Abdul-Aziz et al. 2015; Hansen et al. 2020). This approach is likely beneficial if the population is composed of only drug-susceptible cells. However, if the pathogen population already contains drug-resistant cells, then aggressive treatment may promote the proliferation of the resistant population by eliminating susceptible competitors. To address this problem, an alternative treatment strategy was recently proposed (Day and Read 2016; Hansen et al. 2020). Given that resistance often imposes a cost, resistant variants might be at a competitive disadvantage relative to their sensitive counterparts at low antibiotic concentrations that nonetheless reduce the growth rate of both types. If so, the resulting competition might slow the resistant population’s expansion long enough for the immune system to clear the infection.

Both mathematical models (Hansen et al. 2017) and experiments with the LTEE ancestor (Hansen et al. 2020) have shown that competition between susceptible and resistant populations, mediated in part by fitness costs, can indeed slow the time to treatment failure. However, these expectations are complicated by (i) the potential for higher mutation rates, and (ii) idiosyncratic fitness costs that depend on the specific resistance mutation and its interaction with the genetic background in which it occurs. Regarding the first complication, Hansen and colleagues (2020) used a strain with a low mutation rate (Sniegowski et al. 1997). However, six LTEE populations evolved hypermutability by generation 50,000 (Tenaillon et al. 2016), and mutation rates vary in some pathogens by orders of magnitude (Hughes and Andersson 2017). With respect to the second complication, the competitive release of a resistant population should occur faster when fitness costs are lower. Given that the cost may depend on the particular mutation and its genetic background, the time to treatment failure is harder to predict. We think that these issues and their relevance for treatment options are important avenues for future research.

## Acknowledgments

We thank Chris Adami, Jeff Barrick, Frances Downes, Joshua Franklin, and Chris Waters for valuable discussions and advice, and Devin Lake for general assistance in the laboratory. We acknowledge financial support from an HHMI Gilliam Fellowship (to K.J.C.); a Ralph Evans Award from the MSU Department of Microbiology and Molecular Genetics (to K.J.C.); an NSF grant (currently DEB-1951307 to R.E.L.); a USDA Hatch grant (MICL02253 to R.E.L.); and the BEACON Center for the Study of Evolution in Action (NSF cooperative agreement DBI-0939454).

---

**Table S1.**
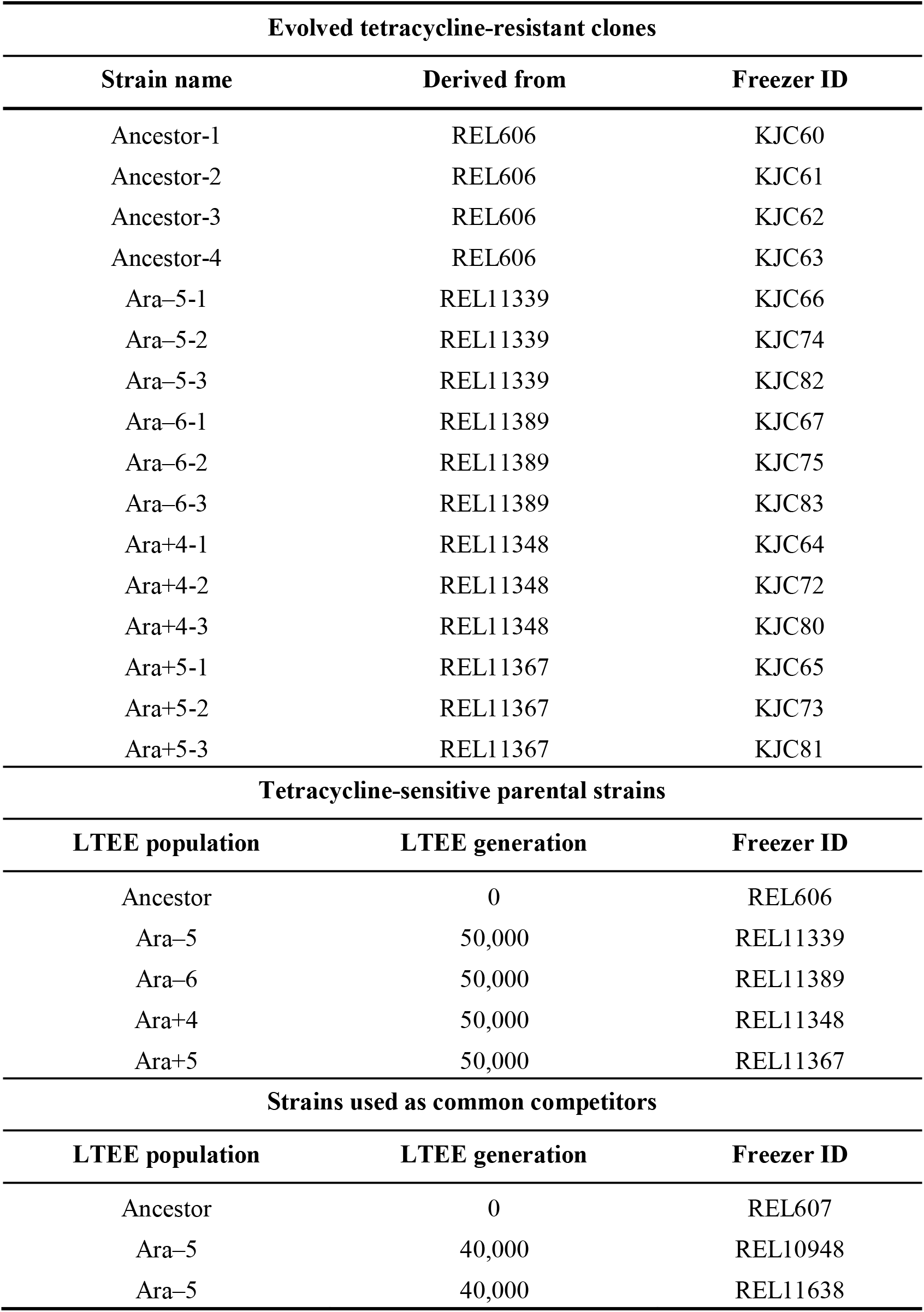
Bacterial strains used in this study.

